# A reusable benchmark of brain-age prediction from M/EEG resting-state signals

**DOI:** 10.1101/2021.12.14.472691

**Authors:** Denis A. Engemann, Apolline Mellot, Richard Höchenberger, Hubert Banville, David Sabbagh, Lukas Gemein, Tonio Ball, Alexandre Gramfort

## Abstract

Population-level modeling can define quantitative measures of individual aging by applying machine learning to large volumes of brain images. These measures of brain age, obtained from the general population, helped characterize disease severity in neurological populations, improving estimates of diagnosis or prognosis. Magnetoencephalography (MEG) and Electroencephalography (EEG) have the potential to further generalize this approach towards prevention and public health by enabling assessments of brain health at large scales in socioeconomically diverse environments. However, more research is needed to define methods that can handle the complexity and diversity of M/EEG signals across diverse real-world contexts. To catalyse this effort, here we propose reusable benchmarks of competing machine learning approaches for brain age modeling. We benchmarked popular classical machine learning pipelines and deep learning architectures previously used for pathology decoding or brain age estimation in 4 international M/EEG cohorts from diverse countries and cultural contexts, including recordings from more than 2500 participants. Our benchmarks were built on top of the M/EEG adaptations of the BIDS standard, providing tools that can be applied with minimal modification on any M/EEG dataset provided in the BIDS format. Our results suggest that, regardless of whether classical machine learning or deep learning was used, the highest performance was reached by pipelines and architectures involving spatially aware representations of the M/EEG signals, leading to R^2 scores between 0.60-0.71. Hand-crafted features paired with random forest regression provided robust benchmarks even in situations in which other approaches failed. Taken together, this set of benchmarks, accompanied by open-source software and high-level Python scripts, can serve as a starting point and quantitative reference for future efforts at developing M/EEG-based measures of brain aging. The generality of the approach renders this benchmark reusable for other related objectives such as modeling specific cognitive variables or clinical endpoints.

**Highlights:**

- We provide systematic reusable benchmarks for brain age from M/EEG signals
- The benchmarks were carried out on M/EEG from four countries > 2500 recordings
- We compared machine learning pipelines capable of handling the non-linear regression task of relating biomedical outcomes to M/EEG dynamics, based on classical machine learning and deep learning
- Next to data-driven methods we benchmarked template-based source localization as a practical tool for generating features less affected by electromagnetic field spread
- The benchmarks are built on top of the MNE ecosystem and the braindecode package and can be applied on any M/EEG dataset presented in the BIDS format

## Introduction

Aging-related disorders of the central nervous system affect hundreds of millions of patients, their caregivers and national health services. Over the past decades, important progress has been made in clinical neuroscience, resulting in improvements to clinical diagnosis and treatment (Walhovd et al. 2010; Ewers et al. 2011). Backed by increasingly advanced analytical methods, this has enabled fine-grained characterization of neurodegenerative conditions (Gaubert et al. 2019; Schumacher et al. 2021; Güntekin et al. 2021). Yet, from a public-health perspective, rather than focusing on pathology, it is essential to detect risk factors early within the general population in order to provide actionable feedback for preventive medicine, e.g., by targeting life-style changes. Such predictions are still challenging. Could it be helpful to look at biological rather than chronological age to better estimate the risk of declining brain health?

Recently, brain age has emerged as a concept for estimating biological aging in the general population (James H. Cole and Franke 2017; Liem et al. 2017; Dosenbach et al. 2010). Biological aging can be inferred from the genome via telomere length, mitochondrial function, epigenetics and other cellular features (Ferrucci et al. 2020; Mather et al. 2011). Yet, the age of a person is only a noisy measure of these cellular processes (people of the same chronological age can have different biological ages). At the same time, biological aging affects brain structure and function (K. S. King et al. 2014), inducing loss of brain volume (Driscoll et al. 2009; Scahill et al. 2003) and characteristic changes in neuronal activity (Cabeza et al. 2002; Damoiseaux et al. 2008; Babiloni et al. 2006). A proxy of biological aging can, thus, be obtained by mapping chronological age to brain data from large populations of subjects using machine learning (Liem et al. 2017; Dadi et al. 2021). The resulting models can be used to compute an expectation of a person’s age given her brain data. This is achieved by quantitatively comparing that person’s brain data to the distribution of brain data across different ages within the general population. This statistical expectation can tell how old (or young) a brain “looks” (Spiegelhalter 2016), hence, predicting the risk of neurological complications potentially more precisely than the chronological age.

This empirical measure of biological aging derived from the general population has proven a useful marker of neurodegeneration and cognitive decline in clinical populations (Cole et al. 2018; Raffel et al. 2017; Denissen et al. 2021; Gonneaud et al. 2021). In these cohorts, patients typically appear to have older brains than their chronological age would suggest. Importantly, similar trends emerge when evaluating brain age in the general population where elevated brain age, compared to chronological age, has been associated with lower cognitive capacity, well-being, and general health (Dadi et al. 2021; Cole 2020; Wrigglesworth et al. 2021). Yet, so far, this approach has mainly been based on anatomical brain scans and hemodynamic signals obtained from magnetic resonance imaging (MRI). This limits the broad utility of brain age for public health, as cerebral MRI scans are usually collected when there is an indication, which can be too late. Even when people from the general population are motivated to participate in brain research, this only concerns a small fraction of society: MRI devices and neuroscientific studies are not equally accessible in all regions of the world and do not attract all people equally from within society, potentially leading to selection bias (Fry et al. 2017).

New hope to generalize this approach has been sparked by advances in large-scale modeling of biomedical outcomes from non-invasive electrophysiological data including magnetoencephalography (MEG) and electroencephalography (EEG) (Gaubert et al. 2019; Engemann et al. 2018). This line of research in clinical neurology may help develop assessments of brain health in many additional contexts in which MRI cannot be applied. First MEG-based brain-age models have allowed to validate MEG-derived brain age against MRI-derived brain age. Results from several studies have shown that the MEG- and MRI-derived brain age are statistically related, leading to overlapping correlations between ensuing brain age estimates (Engemann et al. 2020; Sabbagh et al. 2020; Xifra-Porxas et al. 2021) and individual differences in cognition and health. This overlap can be explained by electromagnetic field spread, independently of neuronal activity: As brain structure changes due to aging, cortical activity, even if unchanged, will project differently onto the M/EEG sensor array, making age indirectly decodable (Sabbagh et al. 2020). Importantly, multiple articles have found that neuronal activity captured by MEG adds specific information not present in MRI-derived brain age (Engemann et al. 2020; Xifra-Porxas et al. 2021), leading to improved prediction performance and richer neurocognitive characterization (Engemann et al. 2020). While MEG can provide an important discovery context, it is unlikely to be the right instrument for addressing the availability issues of MRI-based brain age as MEG scanners are even rarer than MRI scanners. In this context, EEG can make a true difference as EEG is economical and allows for flexible instrumentation for neural assessments in a wide range of clinical and real-world situations including at-home assessments. First evidence suggests that MEG-based strategies for brain-age modeling can be translated to EEG. In an earlier publication (Engemann et al. 2020) we found that among many alternative features of varying data-processing complexity, the spatial distribution of cortical power spectra in the beta (13-30Hz) and alpha (8-13Hz) frequency band explained most of the MEG’s performance as brain-age regressor. This type of information can be well accessed without source localization from the sensor-space covariance using spatial filtering approaches or Riemannian geometry (Sabbagh et al. 2020; D. Sabbagh et al. 2019), which has led to successful translation of this MEG-derived strategy to clinical EEG with around 20 electrodes (David Sabbagh et al. 2020). In clinical and real-world contexts in which EEG is frequently collected, fine-grained spatial information may not be present as only a few electrodes are used. This has favored alternative EEG-derived brain-age models focusing on a wealth of spectral and temporal features (Al Zoubi et al. 2018) which may perform better on sparse EEG-montages and has enabled sleep-based brain age measures (Sun et al. 2019; Ye et al. 2020).

These results provide a sense of the flexibility and future potential of EEG-based brain age as a widely applicable real-world measure of brain health. Yet, to fully develop this research program, more and richer evidence is desirable. At this point, comparisons between different machine learning strategies are difficult. Most models were not only developed and validated in one specific context, but their implementations and data-processing routines are dataset-specific. Moreover, general machine learning approaches successful at pathology decoding should be well-suited for brain age modeling too, yet they have never been tested for that purpose (Gemein et al. 2020; Banville et al. 2020; Engemann et al. 2018). This makes it hard to know whether any strategy is globally optimal and where specific strategies have their preferred niche. As a result, uncertainty is added to comparisons between MEG, EEG and MRI, slowing down efforts of validating M/EEG-based brain age. Finally, to mitigate the impact of selection bias concerning the subjects investigated, it will be crucial to analyze many, socially and culturally diverse M/EEG datasets and find representations that are invariant to confounding effects that can raise issues of fairness and racial bias if remaining unaddressed (Choy, Baker, and Stavropoulos 2021). To develop the next generation of M/EEG-derived brain age models, to facilitate processing of larger numbers of diverse M/EEG-data resources and to avoid fragmentation of research efforts, standardized software and reusable benchmarks are needed.

In this paper we wish to make a first step in that direction. We provide reusable brain-age-prediction benchmarks for different machine learning strategies validated on multiple M/EEG datasets from different countries. The benchmarks are built on top of highly standardized dataset-agnostic code enabled by the BIDS standard (Gorgolewski et al. 2016; Niso et al. 2018; Appelhoff et al. 2019). This makes the benchmarks easy to extend in the future for additional datasets. The paper is organized as follows. The method section motivates the choice of the different machine learning benchmarks. The general data processing approach and software developed for this contribution are presented in the context of the benchmark. The selection of datasets is motivated, and datasets are then described in detail and compared regarding key figures that could provoke differences between benchmarks. Dataset-specific processing steps and peculiarities are highlighted. Then a model validation strategy is developed. The results section presents benchmarks on prediction performance across machine learning models and datasets and different performance metrics. The discussion inspects differences between models, modalities, and datasets, identifying unique niches, safe bets as well as unresolved challenges. The work concludes with practical suggestions on additional benchmarks that can be readily explored using the proposed tools and resources. The scripts and library code for this benchmark are publicly available on GithHub^1^.

## Methods

### Brain age benchmarks

Many different approaches exist for ML in neuroscience, and it can be hard to select among them. The following categorization may help orient practical reasoning and study design. What varies in the taxonomy of methods discussed below is how much M/EEG data are statistically summarized before being presented to the learning algorithm. In other words, ML methods vary with respect to the extent to which compression and summary of the M/EEG signals is performed by the learning algorithm vs. feature-defining procedures performed before and independently of the machine learning algorithm.

#### A-priori defined, a.k.a. handcrafted, features

The first category represents approaches in which features are inspired by theoretical and empirical results in neuroscience or neural engineering. Here, M/EEG is summarized in a rigid fashion by global aggregation across sensors, time, and frequencies or by visiting specific regions of interest (Gemein et al. 2020; Sitt et al. 2014; Engemann et al. 2018). A meaningful composition of features requires prior knowledge of the (clinical) neuroscience literature, especially when interpretation of the model is a priority. In practice, it is convenient to extract all or the most relevant features discussed in a given field, apply multiple spatial and temporal aggregation strategies, and then bet on the capacity of the learning algorithm to ignore irrelevant features (Sitt et al. 2014). This motivates the use of tree-based algorithms like random forests (Breiman 2001) that are easy to tune, can fit nonlinear functions (higher-order interaction effects), and are relatively robust to the presence of uninformative features. As local methods that can be seen as adaptive nearest neighbors (Hastie et al. 2005), the predictions of random forests and related methods are bounded by the minimum and maximum of the outcome in the training distribution. For clinical neuroscience applications, this has proven to yield robust off-the-shelf prediction models that are relatively unaffected by noise in the data and in the outcome (Engemann et al. 2018). This approach is also a natural choice when using sparse EEG-montages with few electrodes.

Here we implemented a strategy pursued in (Gemein et al. 2020) and (Banville et al. 2020), aiming at a broad set of different summary statistics of the time-series or the power spectrum. This approach has turned out useful for a pathology detection task in which the labeling of EEG as pathological can be due to different clinical reasons, hence, affecting many different EEG signatures in potentially diffuse ways. Features were computed using the MNE-features package (Schiratti, Le Douget, Van Quyen, et al. 2018). More specifically we used as features (each computed for individual channels and concatenated across channels, and then averaged across epochs): the standard-deviation, the kurtosis, the skewness, the different quantiles (10%, 25%, 75%, 90%), the peak-to-peak amplitude, the mean, the power ratios in dB among all frequency bands (0 to 2Hz, 2 to 4Hz, 4 to 8Hz, 8 to 13Hz, 13 to 18Hz, 18 to 24Hz, 24 to 30Hz and 30Hz to 49Hz), the spectral entropy (Inouye et al. 1991), the approximate and sample entropy (Richman and Moorman 2000), the temporal complexity (Roberts, Penny, and Rezek 1999), the Hurst exponent as used in (Devarajan et al. 2014), the Hjorth complexity and mobility as used in (Päivinen et al. 2005), the line length (Esteller, Echauz, et al. 2001), the energy of wavelet decomposition coefficients as proposed in (Teixeira et al. 2011), the Higuchi fractal dimension as used in (Esteller, Vachtsevanos, et al. 2001), the number of zero crossings and the SVD Fisher Information (per channel) (Roberts, Penny, and Rezek 1999).

#### Covariance-based filterbank approaches

This category represents approaches in which the spatial dimension of M/EEG is fully exposed to the model, whereas temporal or spectral aspects of the signal are to some extent summarized before modeling. As M/EEG signals reflect linear superposition of neuronal activity projected to the sensors through linear field/potential spread, it is natural to use linear (additive) models for adaptively summarizing the spatial dimension of M/EEG signals (King et al. 2018; Stokes, Wolff, and Spaak 2015; King and Dehaene 2014). This intuition is driving the success of linear decoders for evoked response analysis but faces additional challenges when applied to power spectra (Sabbagh et al. 2020). Computing power features on M/EEG sensor-space signals renders the regression task a non-linear problem for which linear models will provide sub-optimal results (Sabbagh et al. 2019). In practice, this can be overcome by extracting nonlinear features like spectral power after anatomy-based source localization, or in a data-driven fashion that does not require availability of individual MRI scans. Spatial filtering techniques provide unmixing of brain sources based on statistical criteria without using explicit anatomical information, which has led to supervised spatial filtering pipelines (de Cheveigné and Parra 2014; Dähne et al. 2014). Another related strategy consists in computing features that are invariant to field spread. This can be achieved by Riemannian geometry, an approach first applied to M/EEG in the context of brain computer interfaces but that has also proven effective for biomarker learning (Barachant et al. 2012; Yger, Berar, and Lotte 2017; Rodrigues, Jutten, and Congedo 2019). These approaches have in common to favor the covariance of M/EEG sensors as a practical representation of the signals. Manipulating the covariance allows one to suppress the effects of linear mixing while, at the same time, exposing the power spectrum and the spatial structure of neuronal activity in each frequency band (Sabbagh et al. 2020). To scan along the entire power spectrum, one computes covariances from several narrow-band signals covering low to high frequencies (Sabbagh et al. 2020). This provides spatially fine-grained information of frequency-specific neuronal activity, hence the term *filterbank*.

Here we implemented the filterbank models from (Sabbagh et al. 2020; Sabbagh et al. 2019) based on Riemannian geometry that were found to provide a practical alternative to MRI-based source localization, although falling slightly behind in terms of performance. This may be explained by the model violations arising from computing the Riemannian embedding across multiple participants. The Riemannian embedding assumes linear field spread but each recording comes from a different head and different sensor locations, which is explicitly modeled when computing individual-specific source estimates. It is an open question whether template-based source localization can improve upon the Riemannian pipeline, observing that in the case of MEG such a procedure would be informed by the head position in the MEG dewar. Both average brain templates and Riemannian embeddings mitigate field spread in a global way with the difference that the average template uses some anatomical information and approximate sensor locations in the context of MEG, whereas Riemannian embeddings are purely a data-driven procedure with some whitening based on the average covariance (across subjects).

To evaluate the benefit of a template-based anatomy, we included a filterbank model using source localization based on the *fsaverage* subject from FreeSurfer (Fischl 2012). The forward model was computed with a 3-layer Boundary Element Method (BEM) model. Source spaces were equipped with a set of 4098 candidate dipole locations per hemisphere. Source points closer than 5mm from the inner skull surface were excluded. The noise covariance matrices used along with forward solutions to compute minimum-norm estimates inverse operators were taken as data-independent diagonal matrices. Diagonal values defaulted to the M/EEG-specific expected scale of noise (obtained via the “make_ad_hoc_cov” function from MNE-Python). All computations were done with MNE (Gramfort et al. 2014, 2013). For computational efficiency, source power estimates were obtained by applying the inverse operators to the subjects’ covariance data (MNE-Python function “apply_inverse_cov”). Dimensionality reduction was carried out with a parcellation containing 448 ROIs (Khan et al. 2018). This procedure closely followed the one from (Engemann et al. 2020), with the difference that here an MRI template was used instead of subject-specific MRIs. Finally, the 448 ROI-wise source power estimates represented as diagonal matrices were the inputs of the log-diag pipeline from (Sabbagh et al. 2020; D. Sabbagh et al. 2019). Features were computed using the coffeine package^2^.

#### Deep learning approaches

This category concerns modeling strategies in which the outcome is mapped directly from the raw signals without employing separate a priori feature-defining procedures. Instead, multiple layers of nonlinear but parametric transformations are estimated end-to-end to successively summarize and compress the input data. This process is controlled by supervision and enabled by a coherent single optimization objective. In many fields, emerging deep learning methods keep defining the state of the art in generalization performance, often outperforming humans. Deep learning models are however greedy for data, and it may take hundreds of thousands if not millions of training examples until these models show a decisive advantage over classical machine-learning pipelines. Applied to neuroscience, where the bulk of datasets is small to medium-sized, deep learning models may or may not outperform classical approaches (Poldrack, Huckins, and Varoquaux 2020; Schulz et al. 2020; Roy et al. 2019; He et al. 2020). The success of using a deep-learning model may, eventually, depend on the amount of energy and resources invested in its development (Gemein et al. 2020).

Apart from high performance on standard laboratory M/EEG datasets and decoding tasks, deep learning models are attractive for other reasons. First, when very specific hypotheses about data generators or noise generators are available (Kietzmann, McClure, and Kriegeskorte 2019). In this setting, the model architecture can be designed to implement this knowledge, e.g. to explicitly extract band power features in a motor decoding task. Second, these models have a strategic advantage when the data generating mechanism is not known at all, hence, few hypotheses about classes of features are available (Schirrmeister et al. 2017). In this setting, models with a generic architecture can learn and identify relevant features themselves without requiring expert knowledge of the researcher. With neural architecture search and automated hyperparameter optimization, there is also intense research to even reduce the amount of expert knowledge needed to create the network architecture itself. This flexibility has led neuroscientists to discover the framework as a vector for hypothesis-driven research probing brain functions and neural computation (Yamins and DiCarlo 2016; Bao et al. 2020). At the same time, this flexibility is equally beneficial under complex environmental conditions that degrade the quality of M/EEG recordings (e.g. real-world recordings outside of controlled laboratory conditions), in which the classes of relevant features are not a priori known and deep learning models can exploit the structure of the data and noise sources to provide robust predictions. (Banville et al. 2021).

Based on prior work, here we benchmarked two battle-tested general architectures (Gemein et al. 2020) implemented using the Braindecode package^3^ (Schirrmeister et al. 2017; Gramfort et al. 2013). Braindecode is an open-source library for end-to-end learning on EEG signals. It is closely intertwined with other libraries. One of them is Mother of all BCI Benchmarks (MOABB) (Jayaram and Barachant 2018), which allows for convenient EEG-data fetching, MNE (Gramfort et al. 2013, 2014), implements well established data structures, preprocessing functionality, and more. A second key dependency is Skorch (Tietz et al. 2017), which implements the commonly known scikit-learn (Pedregosa et al. 2011) API for neural network training (Buitinck et al. 2013). For these reasons, Braindecode is equally useful for EEG researchers who desire to apply deep learning as well as for deep learning researchers who desire to work with EEG data. Braindecode builds on PyTorch (Paszke et al. 2019) and comprises a zoo of decoding models that were already successfully applied to a wide variety of EEG decoding classification and regression tasks, such as motor (imagery) decoding (Schirrmeister et al. 2017; Kostas and Rudzicz 2020), pathology decoding (Gemein et al. 2020; van Leeuwen et al. 2019; Tibor Schirrmeister et al. 2017), error decoding (Völker et al. 2018), sleep staging (Chambon et al. 2018; Perslev et al. 2021), and relative positioning (Banville et al. 2020).

For this benchmark and the task of age regression we used two Convolutional Neural Networks (ConvNets, sometimes abbreviated CNNs) (LeCun et al. 1999) namely ShallowFBCSPNet (BD-Shallow) and Deep4Net (BD-Deep) (Schirrmeister et al. 2017). BD-Shallow was inspired by the famous filter bank common spatial pattern (FBCSP) (Ang et al. 2008) algorithm. Initially, it has two layers that represent a temporal convolution as well as a spatial filter. Together with a squaring and logarithmic non-linearity it was designed to specifically extract bandpower features. Of note, in the present context this architecture is closely related to SPoC (Dähne et al 2014) and, in therefore, in principle, has the capacity to deliver consistent regression models as was formally proven in previous work (Sabbagh et al 2020).

In contrast, BD-Deep is a much more generic architecture. In total, it has four blocks of convolution-max-pooling and is therefore not restricted to any specific features. While BD-Deep has around 276k trainable parameters and has therefore more learning capacity, BD-Shallow has only about 36k parameters.

It is important to note, that we did neither adjust the model architectures (apart from those changes required by the regression task) nor run task-specific hyperparameter optimization. Both ConvNets were used as implemented in Braindecode with hyperparameters that were already successfully applied to pathology decoding from the TUH Abnormal EEG Corpus (Gemein et al. 2020; van Leeuwen et al. 2019; Tibor Schirrmeister et al. 2017). For more information on Braindecode or the ConvNets, please refer to the original publication (Schirrmeister et al. 2017). For decoding, we converted the MEG input data from Tesla to Femtotesla, the EEG input data from Volts to Microvolts, and additionally rescaled the data, such that it has roughly zero mean and unit variance by dividing by the standard deviation of each dataset (see Section Datasets).

### General data processing strategy using BIDS and the MNE-BIDS pipeline

Neuroimaging and behavioral data are stored in many different complex formats, potentially hampering efforts of building widely usable methods, hence, impeding reproducible research. Our goal was to provide brain-age prediction models that can be directly applicable to any new electrophysiological dataset. For this purpose, we used the Brain Imaging Data Structure (BIDS) (Gorgolewski et al. 2016) which allows us to organize neuroimaging data in a standardized way supporting interoperability between programming languages and software tools. We used the MNE-BIDS software (Appelhoff et al. 2019) for programmatically converting M/EEG datasets into the BIDS format (Pernet et al. 2019; Niso et al. 2018). This has allowed us to access all datasets included in this work in the same way, enabling data analysis for all these datasets with the same code. We will now summarize the general workflow (cf. *Fig. 1)*.

**Figure 1:**
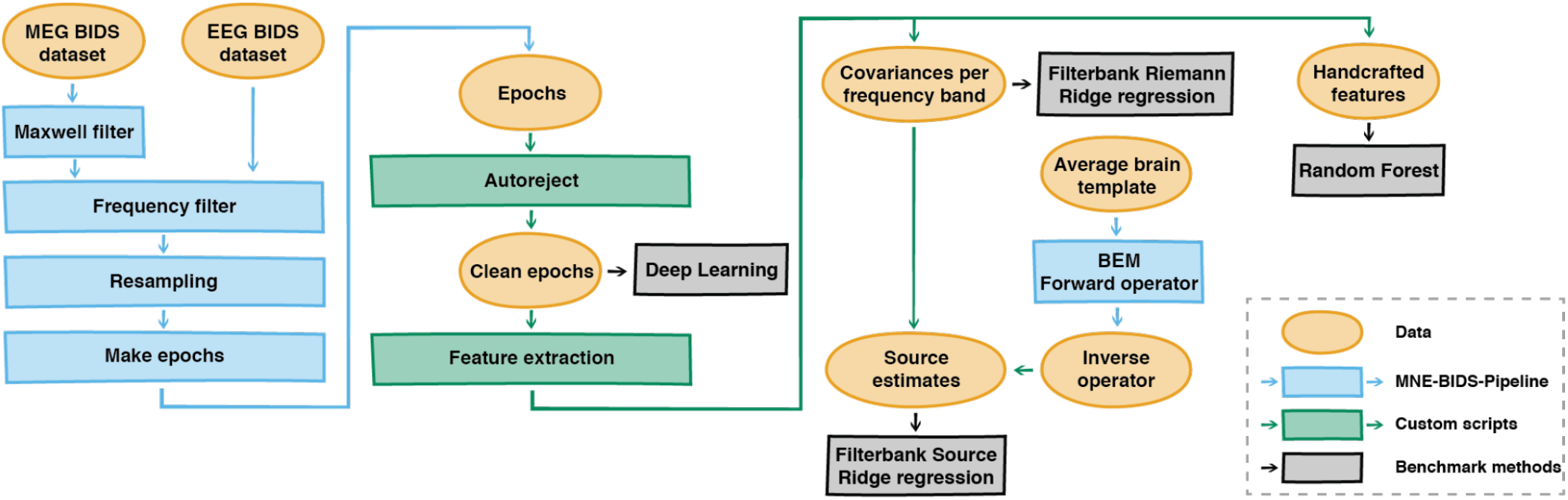
Data processing, feature extraction and model construction based on the BIDS standard. This benchmark project provides a common data processing and feature extraction code allowing comparisons of different classical and deep learning-based machine learning models across different M/EEG datasets. Support for new datasets can be added with minimal modifications. For a detailed description consider the main text and the open-source code repository supporting this article^5^.

For this study, we used the MNE-BIDS-Pipeline for automatic preprocessing of MEG and EEG data stored in BIDS format^4^ (Jas et al. 2018). Its main advantage is that we can implement various custom analyses for different datasets without having to write any elaborate code. Modifying the overall processing pipeline or adapting a given pipeline to a new dataset only requires few edits. Controlling the pipeline is achieved through dataset-specific configuration files that specify the desired processing steps and options of the MNE-BIDS-Pipeline while dealing with the peculiarities of the data. The MNE-BIDS-Pipeline scripts themselves do not need to be modified and are readily applicable on diverse datasets.

We designed configuration files to implement data processing steps common to all datasets analyzed in this benchmark while handling dataset-specific details. Raw signals bandpass-filtered between 0.1 and 49Hz using a zero-phase finite impulse response (FIR) filter with Hamming window. Window length and transition bandwidth were automatically controlled by default settings of MNE-Python (v0.24). We considered epochs of 10-second length without overlap. These epochs coincided with eyes-closed or eyes-open resting-state conditions in some of the datasets. As additional channels measuring ocular and cardiac activity were not consistently available across datasets, we only implemented amplitude-based artifact rejection using the local autoreject method (Jas et al. 2017). Through 5-fold cross-validation, autoreject chose channel-specific rejection peak-to-peak-amplitude thresholds and then decided if a given epoch could be repaired using interpolation, or if it should be rejected to obtain clean data. We kept the default grid of candidate values for the hyperparameters ‘rho’ (the consensus proportion of bad channels leading to rejection of an epoch) and ‘kappa’ (maximum number of channels allowed to be interpolated). For ‘rho’ we considered a linearly spaced grid of 11 points between 0 and 1. For ‘kappa’ we considered 1, 4, or 32 channels. As the local autoreject is not yet supported in the MNE-BIDS pipeline, this step was implemented in a custom script (see the “compute_autoreject.py” in the code repository). Apart from preprocessing, we also made use of the MNE-BIDS-Pipeline to generate forward solutions and inverse operators for the source localization approach based on template MRI (see section *Covariance-based filterbank approaches* for detailed explanations).

Each model of the benchmark is based on features extracted from clean epochs. Again, the conversion of datasets to BIDS has enabled feature extraction using one general script for all datasets (“compute_features.py” in the code repository).

### Datasets

Large datasets and biobanks are the backbone of population modeling. In the past 10 years, this has led to a wealth of publications in cognitive neuroscience on modeling biomedical outcomes and individual differences in cognition from MRI data (Kernbach et al. 2018; James H. Cole 2020; Smith et al. 2015). This has been enabled by consortia and large-scale institutional collaborations (Bycroft et al. 2018; Van Essen et al. 2013) that aim at recontextualizing existing data for open-ended future usage (Leonelli 2016). More recently, the first M/EEG datasets have emerged with a focus on characterizing populations (Taylor et al. 2017; Larson-Prior et al. 2013; Babayan et al. 2019; Obeid and Picone 2016; Niso et al. 2016; Valdes-Sosa et al. 2021; Bosch-Bayard et al. 2020). The selection of datasets for the present study did not aim at comprehensiveness but represents an attempt to secure a minimum degree of diversity. Social bias and fairness are important challenges, not only in the field of machine learning but also in biomedical research. It has been shown for modern biobanks that the sample deviates from the general population in important ways, oversampling Caucasian people with higher education degrees (Fry et al. 2017; Henrich and Heine 2010). For deployment of predictive biomarkers, this can have tragic consequences as clinical utility may depend on sex and ethnicity (Duncan et al. 2019). As a result, in EEG research, specific risks of racial bias have been recognized lately, highlighting the risk of selection bias and confounding, e.g., due to culture-specific hair style (Choy, Baker, and Stavropoulos 2021). Taken together, this emphasizes the importance of benchmarking on socially and culturally different datasets. Our selection includes M/EEG datasets from four different countries representing culturally and socioeconomically diverse contexts. In the following we will provide a high-level introduction to the datasets, highlighting characteristic differences, challenges and opportunities for unique benchmarks.

### Cam-CAN MEG data

The Cambridge Centre of Ageing and Neuroscience (Cam-CAN) dataset (Taylor et al. 2017; Shafto et al. 2014) has been the starting point of our efforts in building brain age models (Engemann et al. 2020; David Sabbagh et al. 2020) and we like to see it as a discovery context. The combination of a wide, almost uniformly distributed age range and MEG data alongside MRI and fine-grained neurobehavioral results make it a rich resource for exploring aging-related cortical dynamics. On the other hand, models developed on this dataset may not be generalizable to real-world contexts in which EEG is operated. The following two sections are based on the methods description from our previous publications (Engemann et al. 2020; Sabbagh et al. 2020).

#### Sample description

The present work was based on the latest BIDS release of the Cam-CAN dataset (downloaded February 2021). We included resting-state MEG recordings from 646 participants (female = 319, male = 327). The age of the participants ranged from 18.5 to 88.9 years with a mean age of 54.9 (female = 54.5, male = 55.4) and a standard deviation of 18.4 years. Data is provided in Tesla and has a standard deviation of 369.3 Femtotesla. We did not apply any data exclusion. Final numbers of samples reflect successful preprocessing and feature extraction. For technical details regarding the MEG instrumentation and data acquisition, please consider the reference publications by the Cam-CAN (Taylor et al. 2017; Shafto et al. 2014). In the following we highlight a few points essential for understanding our benchmarks on the Cam-CAN MEG data.

#### Data acquisition and processing

MEG was recorded with a 306 VectorView system (Elekta Neuromag, Helsinki). This system allowed measuring magnetic fields with 102 magnetometers and 204 orthogonal planar gradiometers inside a light magnetically shielded room. During acquisition, an online filter was applied between around 0.03Hz and 1000Hz. After bandpass filtering (0.1 - 49Hz), we applied decimation by a factor of 5, leading to a sample frequency of 200Hz (at the epoching stage). To mitigate the contamination of the MEG signal by environmental magnetic interference, we applied the temporal signal-space-separation (tSSS) method (Taulu, Simola, and Kajola 2005). Default settings were applied for the harmonic decomposition (8 components of the internal sources, 3 for the external sources) on a 10-s sliding window. To discard segments for which inner and outer signal components were poorly distinguishable, we applied a correlation threshold of 98%. As a result of this procedure, the signal was high pass filtered at 0.1Hz and the dimensionality of the data was reduced to 65, approximately. It is worthwhile to note that Maxwell filtering methods like tSSS merge the signal from magnetometers and gradiometers into one common low-rank representation. As a result, after tSSS, the signal displayed on magnetometers becomes a linear transformation of the signals displayed on the gradiometers. This leads to virtually identical results when conducting analyses exclusively on magnetometers versus gradiometers (Garcés et al. 2017). To reduce computation time, we analyzed the magnetometers for our benchmark. To deal with the reduced data rank, a PCA projection to the common rank of 65 was applied whenever the machine learning pipeline was sensitive to the rank (e.g., Riemannian filterbank models). For the full specification of the preprocessing, please refer to the “config_camcan_meg.py” file in the code repository.

### LEMON EEG data

The Leipzig Mind-Brain-Body (LEMON) dataset offers rich multimodal EEG, MRI and fMRI data for a well characterized group of young and elderly adults sampled from the general population (Babayan et al. 2019). As it was the case for the Cam-CAN data, here the research was conducted in a research context using high-end equipment accompanied by rich and fine-grained neurocognitive and behavioral assessments.

#### Sample description

EEG resting-state data from 227 healthy individuals from the LEMON dataset were included in this study. This sample contains 82 females (mean age = 44.2) and 145 males (mean age = 36), representing a clearly visible difference in the composition of the sample (*Fig*. 2). Their age distribution went from 20 to 77 years old with an average of 38.9 +- 20.3 years. Our sample covers the whole available dataset (downloaded September 2021) as we did not apply any exclusion criteria. It is a peculiarity of this dataset is that it is divided into 2 distinct age subpopulations, one between 20-35, the second between 55-77 (*Fig*. 2), rendering the mean a bad representation of the age distribution. Moreover, the public version of the datasets only provides ages in a granularity of 5 years to mitigate the risk of identifying participants. For the purpose of this study, we included the precise ages obtained through institutional collaboration. The impact on average modeling results turned out negligible, however. Data is provided in Volts and has a standard deviation of 9.1 Microvolts.

**Figure 2:**
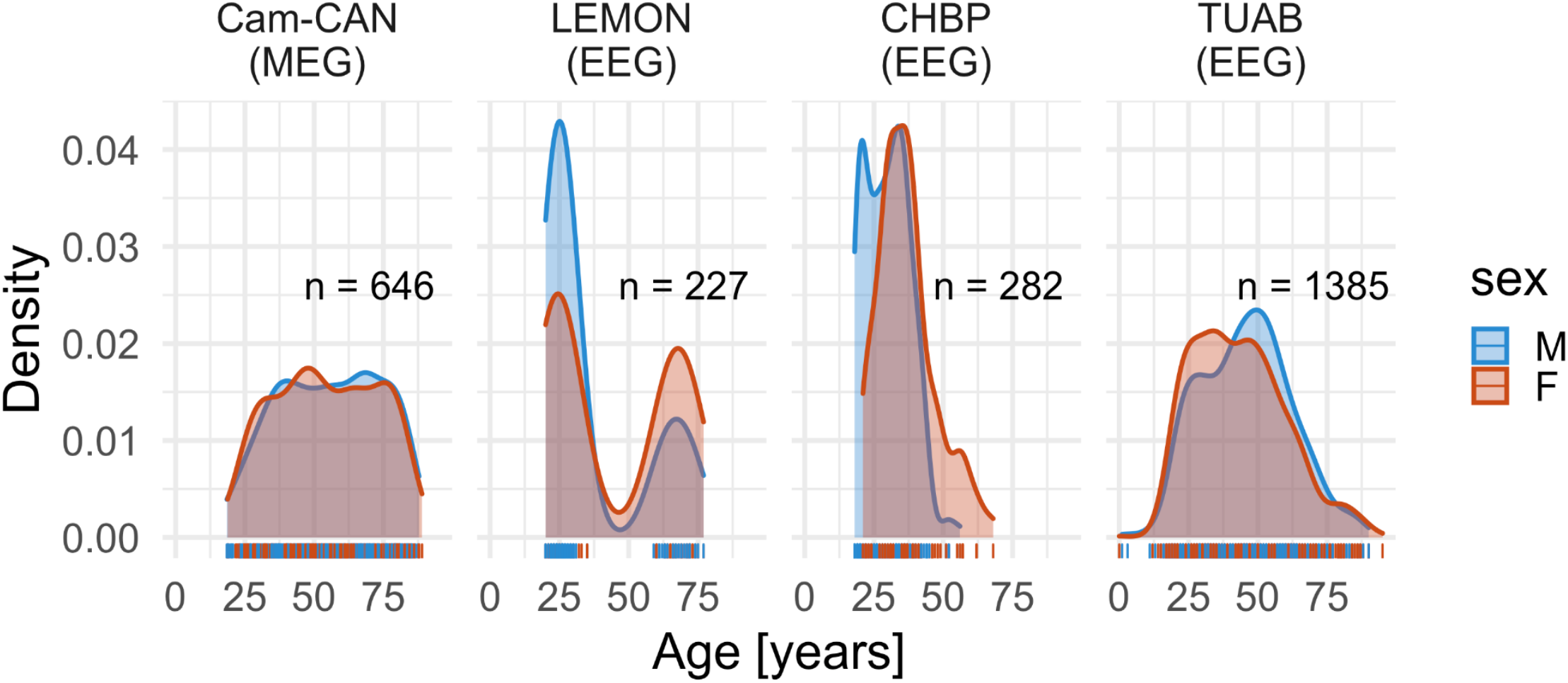
Age distributions by gender by dataset. The kernel density (y axis) is plotted across the age range (x axis) for all four M/EEG datasets included in the study, separately for male (blue) and female (red) participants. Individual observations are displayed by rug plots at the bottom of each panel. The Cam-CAN data (MEG) show a wide age range with a quasi-uniform distribution and no obvious sex imbalance. This situation poses no a priori challenges for age prediction while, at the same time, analysis of MEG data may be more complex. The LEMON dataset included a group of young participants and a group of old participants, leading to a characteristic bi-modal distribution. Sex imbalance is clearly visible with more male participants in the group of young participants and fewer male participants in the group of older participants. This may lead to potential sex differences in prediction success and renders the average age a bad summary of the age distribution. The CHBP data shows a rather reduced age range with a right-skewed age distribution and some sex imbalance (again more young male participants). Predicting the age can be expected to turn out more difficult on this dataset for the implied lack of density along the age range. Finally, the TUAB data present a symmetric age distribution with minor sex differences, however, a less uniform age distribution. This may lead to more pronounced errors in young and elderly participants. This may, however, be compensated for by the more generous sample size. To summarize, the four datasets investigated here pose unique challenges for M/EEG brain age modeling.

#### Data acquisition and processing

EEG was recorded with 62-channel active ActiCAP electrodes and a bandpass filter between 0.015Hz and 1kHz. We applied additional bandpass filtering between 0.1Hz and 49Hz. The channel placement implemented the 10-5 system (Oostenveld and Praamstra 2001). EEG data were sampled at 2500Hz. After bandpass filtering (0.1 - 49Hz), data were decimated by a factor of 5, yielding a final sampling frequency of 500Hz. As a peculiarity of the dataset, resting-state recordings encompass samples from two conditions: eyes-closed and eyes-open. Our pipeline explicitly respected these different conditions. To include a maximum of data and, potentially, a larger set of distinguishable EEG sources, we pooled the data prior to feature extraction. For the full specification of the preprocessing, please refer to the “config_lemon_eeg.py” file in the code repository.

### CHBP EEG data

The Cuban Human Brain Mapping Project (CHBP) provides rich multimodal EEG and MRI data sampled from young to middle-aged adults from the general population (Valdes-Sosa et al. 2021; Hernandez-Gonzalez et al. 2011; Bosch-Bayard et al. 2020). As for the Cam-CAN and LEMON data, research was carried out using high-end electrophysiological equipment in a biomedical research context. However, the data was collected in a Latin American mid-income country, (Valdes-Sosa et al. 2021), adding a much-needed opportunity for increasing the diversity in population-level neuroscience datasets. This diversity expresses itself in the composition of EEG protocols which contain elements of real-world neurology exams, e.g., a hyperventilation task.

#### Sample description

EEG resting-state data from 282 healthy individuals from the CHBP dataset were included in this study. The sample contained 87 females (mean age = 36.7) and 195 males (mean age = 29.9), representing a clearly visible difference in the composition of the sample (*Fig*. 2). The overall age distribution went from 18 to 68 years with an average of 32 +/− 9.3 years. Data is provided in Volts and has a standard deviation of 6.6 Microvolts. Our sample covers the whole available dataset (download June 2021) as we did not apply any exclusion criteria. Final numbers reflect successful processing of the data.

#### Data acquisition and processing

EEG data were recorded using a MEDICID 5 system and two different electrode caps of either 64 or 128 channels. The channel placement implemented the 10-5 system (Oostenveld and Praamstra 2001). Here we focused the analysis on the subset of common channels present in all recordings, leading to 53 channels. We applied additional bandpass filtering between 0.1Hz and 49Hz. As in the LEMON dataset, resting-state recordings encompassed samples from eyes-closed and eyes-open conditions. Again, we pooled both conditions prior to feature extraction. Note that for the data release (downloaded July 2021) used in this work, we could not benefit from the expert-based annotations of clean data. The results obtained on this dataset may therefore be impacted by quality issues to unknown extents.

For the full specification of the preprocessing, please refer to the “config_chbp_eeg.py” file in the code repository.

### TUAB EEG data

The Temple University Hospital Abnormal EEG Corpus (TUAB) provides socially and ethnically heterogeneous clinical EEG data (Obeid and Picone 2016) mostly from Latin-American and African American participants (personal communication, Joseph Picone). As a peculiarity, the EEG data is obtained from an archival effort of recovering different EEG exams from the Temple University Hospital in Philadelphia. The clinical and social diversity render the TUAB dataset an important resource for electrophysiological population modeling (Gemein et al. 2020; David Sabbagh et al. 2020).

#### Sample description

Here, we focused exclusively on the EEG recordings labeled as not pathological by medical experts comprising a subsample of 1385 subjects (female = 775 and males = 610). This sample contained individuals ranging from newborn children (min age = 0 for female and min age = 1 for male) to elderly (max age = 95 for female and 90 for male) people (*Fig*. 2). The average age is 44.4 +/− 16.5 years. Data is provided in Volts and has a standard deviation of 9.7 Microvolts. The data processing closely followed our previous work on the TUAB data (Sabbagh et al. 2020). For further details about the dataset, please refer to the reference publications (Harati et al. 2014; Obeid and Picone 2016).

#### Data acquisition and processing

EEG data were recorded using different Nicolet EEG devices (Natus Medical Inc.), equipped with between 24 and 36 channels. For channel placement, the 10-5 system was applied (Oostenveld and Praamstra 2001). All sessions have been recorded with an average reference. Here we considered a subset of 21 common channels. As channel numbers differed across recordings, re-referencing was necessary. For consistency, we also applied re-referencing with an average reference on all other EEG datasets. As sampling frequencies were inconsistent across recordings, we resampled the data to 200Hz. For many patients, multiple recordings were available. For simplicity we only considered the first recording. For the full specification of the preprocessing, please refer to the “config_tuab_eeg.py” file in the code repository.

### Model evaluation and comparison

To gauge model performance, we first defined a baseline model that should not provide any intelligent prediction. As in previous work (Sabbagh et al. 2020; Sabbagh et al. 2019; Engemann et al. 2020), we employed a dummy regressor model as a low-level baseline in which the outcome is guessed from the average of the outcome on the training data. This approach is fast and typically converges with more computationally demanding procedures based on permutation testing that we shall briefly outline.

This is particularly relevant for the present benchmark where the combinatorial matrix of machine learning models (including deep learning) versus datasets would lead to unpleasant computation times when applying tens of thousands of permutations. The same can be said for other approximations focusing on ranking statistics across hundreds of Monte Carlo cross-validation iterations (Sabbagh et al. 2019). Finally, another approach relies on large left-out datasets, entirely independent from model construction, in which predictions can be treated like random variables, hence, classical inferential statistics are valid. In previous work (Dadi et al. 2021), permutation tests and the non-parametric bootstrap were employed on more than 4000 left-out data points to assess performance above chance and pairwise differences between models. Such generous held-out datasets are not available in the present setting, nor can we readily compute statistics across folds, as cross-validation iterations are not statistically independent. We therefore implemented a less formal approach comparing competing models against dummy regressors and against each other based on standard 10-fold cross-validation based on fixed random seeds. This ensured that for any model under consideration, identical data splits were used. Of note, our reusable benchmark code allows interested readers to implement more exhaustive model comparison strategies.

For scoring prediction performance, we focused on two complementary metrics. The coefficient of determination (R^2^) score and the mean absolute error (MAE). Considering the dummy regressor, the R^2^ score is a natural choice as it quantifies the incremental success of a model over a regressor returning the average of the training-data as a guess for the outcome. Compared to Pearson correlations that are sometimes used in applied neuroscience studies, the R^2^ metric is more rigorous as it is sensitive to the scale of the error and the location: Predictions that are entirely biased, e.g, shifted by a large offset, could still be correlated with the outcome. In contrast, the R^2^ metric clearly penalizes systematically wrong predictions by assigning scores smaller than 0. Positive predictive success thus falls into a range of R^2^ between 0 and 1. This facilitates comparisons across models within the same dataset while posing challenges when comparing models across datasets.

We therefore considered the MAE which has the benefit of expressing prediction errors at the scale of the outcome. This is particularly convenient for scientific interpretation when the outcome has some practical meaning as is the case in the present benchmarks on age prediction. Importantly, the MAE does not per se resolve the problem of comparisons across datasets as the meaning of errors entirely depend on the distribution of the outcome: Small errors in years are good for datasets with wide age distributions but bad in datasets with narrow age distributions. This obviously calls for contextualizing the MAE against a dummy baseline regression model. While this does not necessarily facilitate comparisons across datasets, it helps make visible situations in which one cannot rely solely on the R^2^ for model comparisons.

### Computational considerations and software

#### M/EEG data processing

BIDS conversion and subsequent data analysis steps were carried out in Python 3.7.1, the MNE-Python software (v0.24, Gramfort et al. 2014, 2013), the MNE-BIDS package (v0.9, Appelhoff et al. 2019) and the MNE-BIDS-pipeline on a 48-core Linux high-performance server with 504 GB RAM. The joblib library (v1.0.1) was used for parallel processing. For artifact removal, the latest development version (v0.3dev) of the autoreject package (Jas et al. 2017) was used.

##### Classical machine learning benchmarks

For future computation, the mne-features (0.2, Schiratti, Le Douget, Le Van Quyen, et al. 2018), PyRiemann (v0.2.6) and the coffeine (0.1, Sabbagh et al. 2020) libraries were used. Analyses were composed in custom scripts and library functions based on the Scientific Python Stack with NumPy (v1.19.5, Harris et al. 2020), SciPy (v1.6.3, Virtanen et al. 2020) and pandas (v.1.2.4, McKinney and Others 2011). Machine-learning specific computation was composed using the scikit-learn package (Pedregosa et al. 2011). Analysis was carried out on a 48-core Linux high-performance server with 504 GB RAM. Feature extraction, depending on the dataset, completed within several hours to days. Model training and evaluation completed within a few minutes to hours. However, feature computation could last several days, depending on the dataset and the types of features.

##### Deep learning benchmarks

A high-performance Linux server with 72 cores, 376 GB RAM and 1 or 2 Nvidia Tesla V100 or P4 GPUs was used. Code was implemented using the PyTorch (Paszke et al. 2019) and braindecode (Schirrmeister et al. 2017) packages. Model training and evaluation completed within 2-3 days.

##### Data visualization

Graphical displays and tables were composed on an Apple Silicon M1 Macbook Pro (space gray) in R (v4.0.3 “Bunny-Wunnies Freak Out”) using the ggplot2 (v3.3.5, Wickham 2011), patchwork (v1.1.1, Pedersen 2019), ggthemes (v4.2.4) and scales (v1.1.1, Arnold 2017) packages with their respective dependencies.

## Results

For the age prediction benchmark, we considered five alternative approaches: heterogeneous handcrafted features & random forest (‘handcrafted’), filterbank features based on Riemannian embeddings & ridge regression (‘filterbank-riemann’), filterbank features based on source localization with MRI-average template & ridge regression (‘filterbank-source’), a shallow deep learning architecture (“shallow”) and a 4-layer deep-learning architecture (‘deep’). These approaches were benchmarked across four M/EEG datasets: The Cambridge Centre of Ageing and Neuroscience (Cam-CAN) dataset (Taylor et al. 2017), the Cuban Human Brain Mapping Project (CHBP) dataset (Valdes-Sosa et al. 2021), the Leipzig Mind-Brain-Body (LEMON) dataset (Babayan et al. 2019) and the Temple University Hospital Abnormal EEG Corpus (TUAB) dataset (Obeid and Picone 2016). Generalization performance was estimated using 10-fold cross validation after shuffling the samples (fixed random seed). The coefficient of determination (R^2^) was used as a metric enabling comparisons between datasets independently of the age distribution, mathematically quantifying the additional variance explained by predicting better than the average age. A dummy model empirically quantifies chance-level prediction by returning the average age of the training data as prediction. The results are displayed in Fig. 3. One can see that on most of the datasets all machine learning models achieved R^2^ scores well beyond the dummy baseline. The highest scores were observed on the Cam-CAN MEG dataset, followed by the LEMON EEG dataset. Caution is warranted though to avoid premature conclusions: The R^2^ offers a common scale that explicitly compares the incremental model performance over the average predictor. This is achieved by dividing the sum of squares of the model’s prediction by the sum of squares of the average predictor but, in turn, depends on the distribution of age. As a result, this can be misleading in cross-dataset comparisons when the variance of the outcome is not the same, which is the case here (cf. Fig. 2). We therefore also computed results using the mean absolute error as a performance metric (Fig 4). One can now see that the overall distribution of scores, including the scores of the dummy model, depend not only on the dataset but also on its age range. Where the range is small, improvements over the baseline models are harder to observe. Moreover, comparing MAE scores across datasets without taking into account the baseline can yield misleading conclusions. For example, the same score of *e.g*. an MAE = 10 can be way above chance in one dataset (Cam-CAN) but below chance in another dataset (CHBP). To alleviate this problem, normalized MAE scores have been suggested in which the MAE scores are related to the range of the age distribution (Cole, Franke, and Cherbuin 2019). This does not come without its own problems, as then outliers in non-uniform distributions could drive the scores. As research keeps evolving on this topic and the community has not yet agreed on the best metric, we recommend considering multiple classical machine learning metrics when comparing model performance – in critical awareness of their respective limitations.

**Fig. 3.**
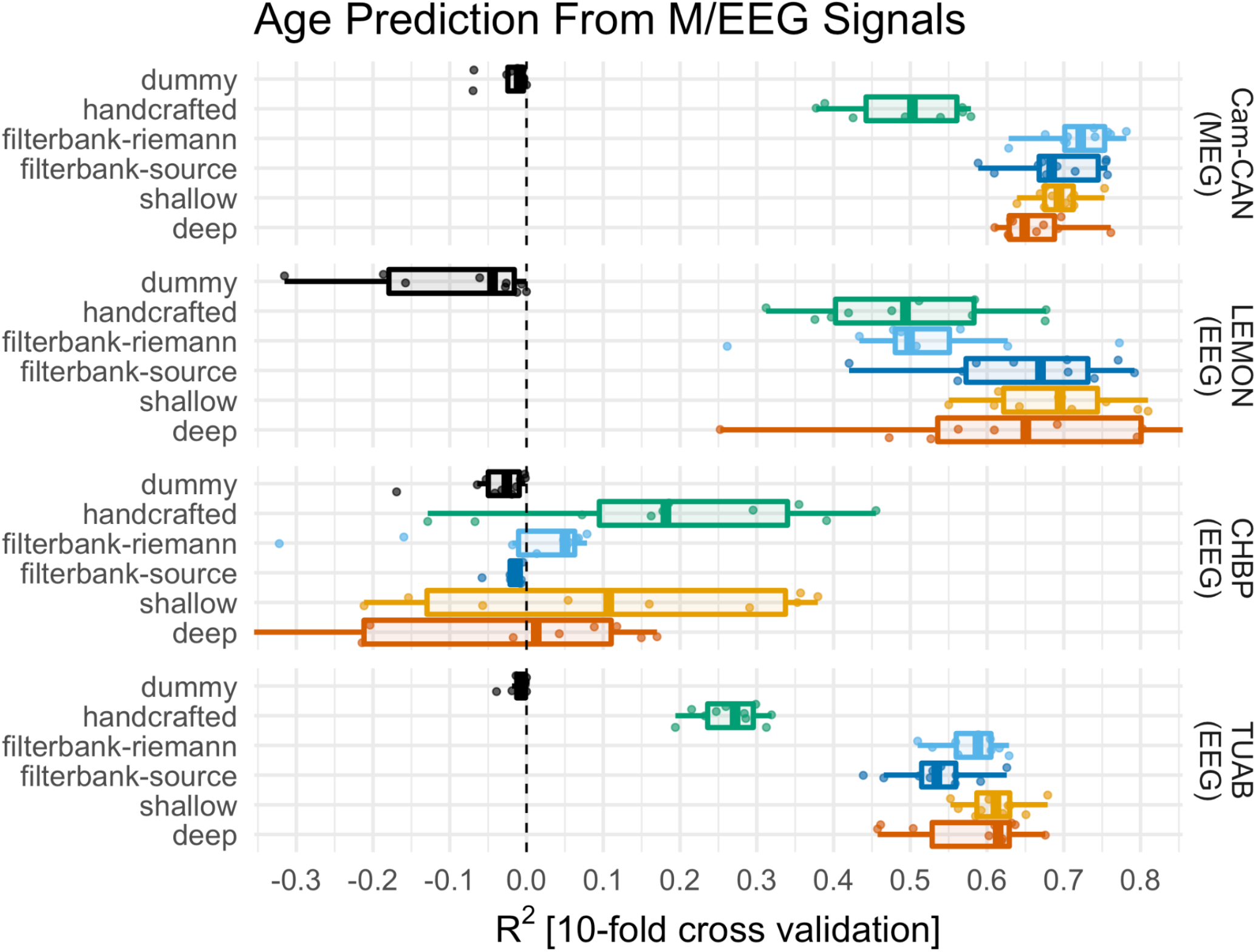
Age prediction benchmarks across M/EEG datasets (R^2^ score). Generalization performance was assessed by 10-fold cross-validation and the R^2^ score for five machine learning strategies compared against a dummy model (rows) and four datasets (panels). Across datasets, dummy models were mostly well-calibrated with R^2^ scores close to zero. The LEMON dataset was one exception as dummy scores were systematically worse than chance, which can be explained by the bimodal age distribution (cf. Fig. 2), rendering the average age a bad guess for the age. The ‘handcrafted’ benchmark delivered moderate but systematic prediction success across all datasets. The two filterbank models performed well across datasets with similar performances, markedly higher than for the ‘handcrafted’ approach. The only exception was the CHBP benchmark for which neither the filterbank nor the deep models delivered useful predictions. Note that here, for the ‘filterbank-source’, a single fold with an abysmal R^2^ score of −15 was obtained (x limits constrained to a range between −.3 and 1.0). Overall, the deep learning benchmarks performed similarly to the filterbank models.

**Fig. 4.**
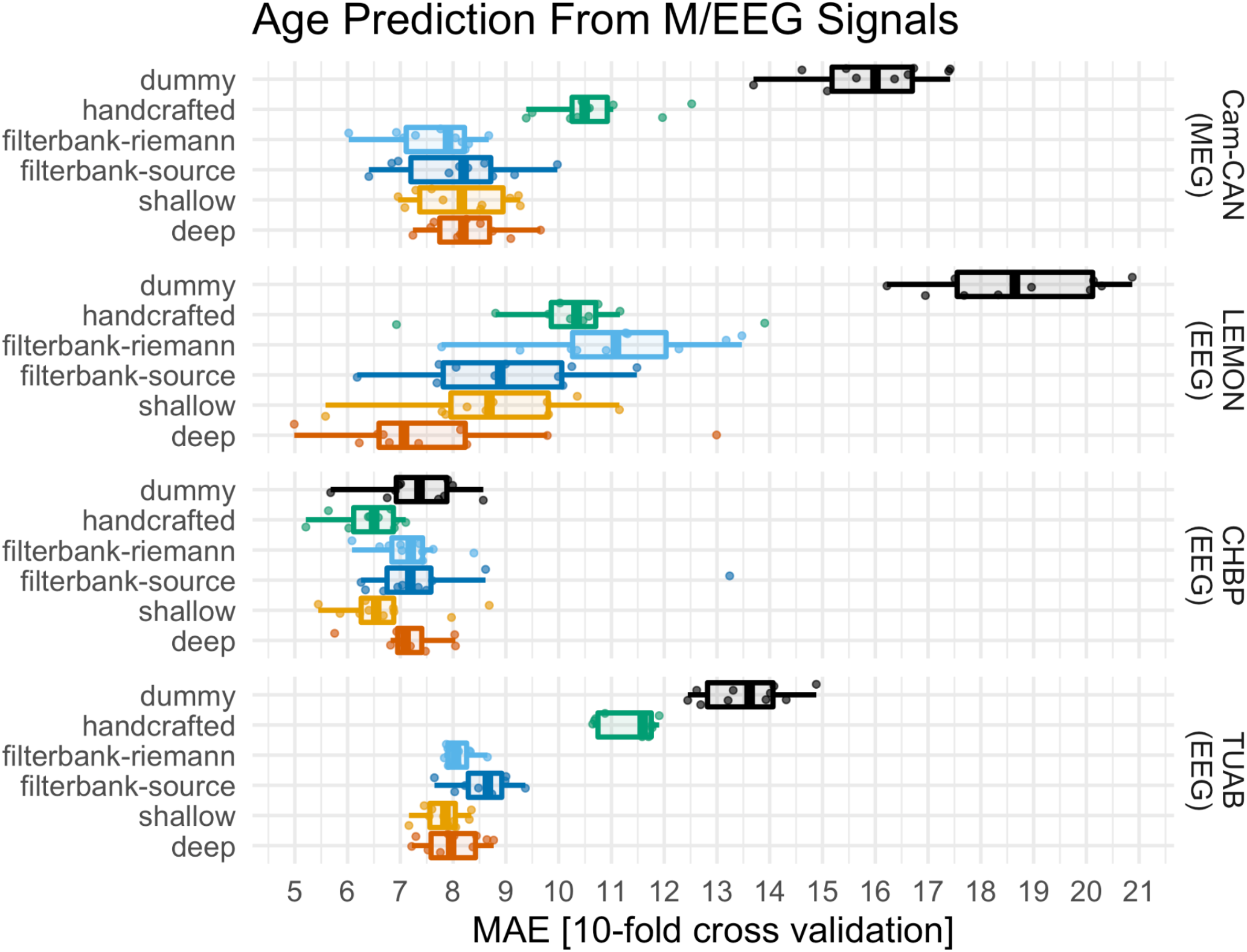
Age prediction benchmarks across M/EEG datasets (mean absolute error). Same visual conventions as in Fig. 3. As the mean absolute error (MAE) is sensitive to the scale and distribution of the outcome, one can see characteristic differences across datasets. The distribution of the dummy scores provides an estimate of the random guessing. As before, in all but the Cuban datasets all benchmarks achieved MAE scores markedly better than the dummy with no overlap between model and dummy distributions. Model rankings resemble the ones obtained using the R^2^. On the LEMON data, the deep benchmark now presented a slight advantage over all other benchmarks.

Confronting the relative performances of models to the dummy baseline in Fig. 3 and Fig. 4, one can see overall similar performance rankings between the models, regardless of the metric. See Table 1 for side-by-side comparisons of the aggregated cross-validation distributions. The big-picture results argue in favor of the importance of fine-grained spatial features for M/EEG prediction while considering important between-dataset heterogeneity. Both filterbank pipelines provide features based on spatially aware representations of the M/EEG signals, which either explicitly or implicitly deal with the spatial spread of electrical potentials and fields characteristic for M/EEG signals. The source-level filterbank approximates source localization using the average MRI template, whereas Riemannian embeddings provide non-linear spectral features that are affine invariant, hence, independent of linear mixing. The deep benchmarks, on the other hand, implied spatial-filtering layers capable of mimicking source localization by learning an unmixing function. Surprisingly, using the average MRI template instead of the Riemannian embedding to construct a filterbank model did not lead to consistent improvements across datasets, suggesting that both approaches may be equally effective in practice. We would have conjectured that even an imprecise biophysical head model would provide inverse solutions leading to more accurate unmixing of M/EEG sources. Compared to our previous benchmarks (Engemann et al. 2020; Sabbagh et al. 2020) favoring filterbank models based on source-localization, one has to point out that this finding may reflect at least two differences: The use of an MRI template instead of individual co-registration and the use of empty-room-based suppression of environmental noise. The second factor may be less relevant for EEG though where empty room recordings are not available and data-based covariances are more common in event-related studies where brain activity induced by stimuli is compared against the background resting-state activity. As a practical implication, and if inspection of the brain sources is not a priority, the purely data-driven pipelines may be more practical as no additional MRI-based data processing is needed (cf. Fig. 1).

**Table 1.**
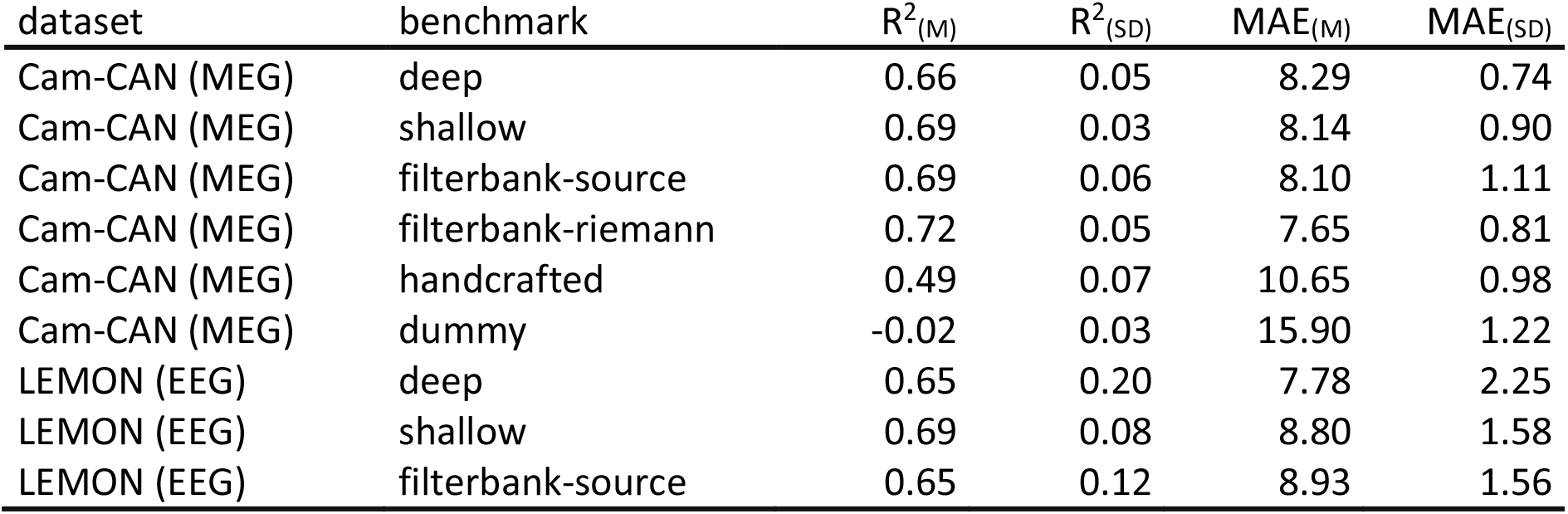
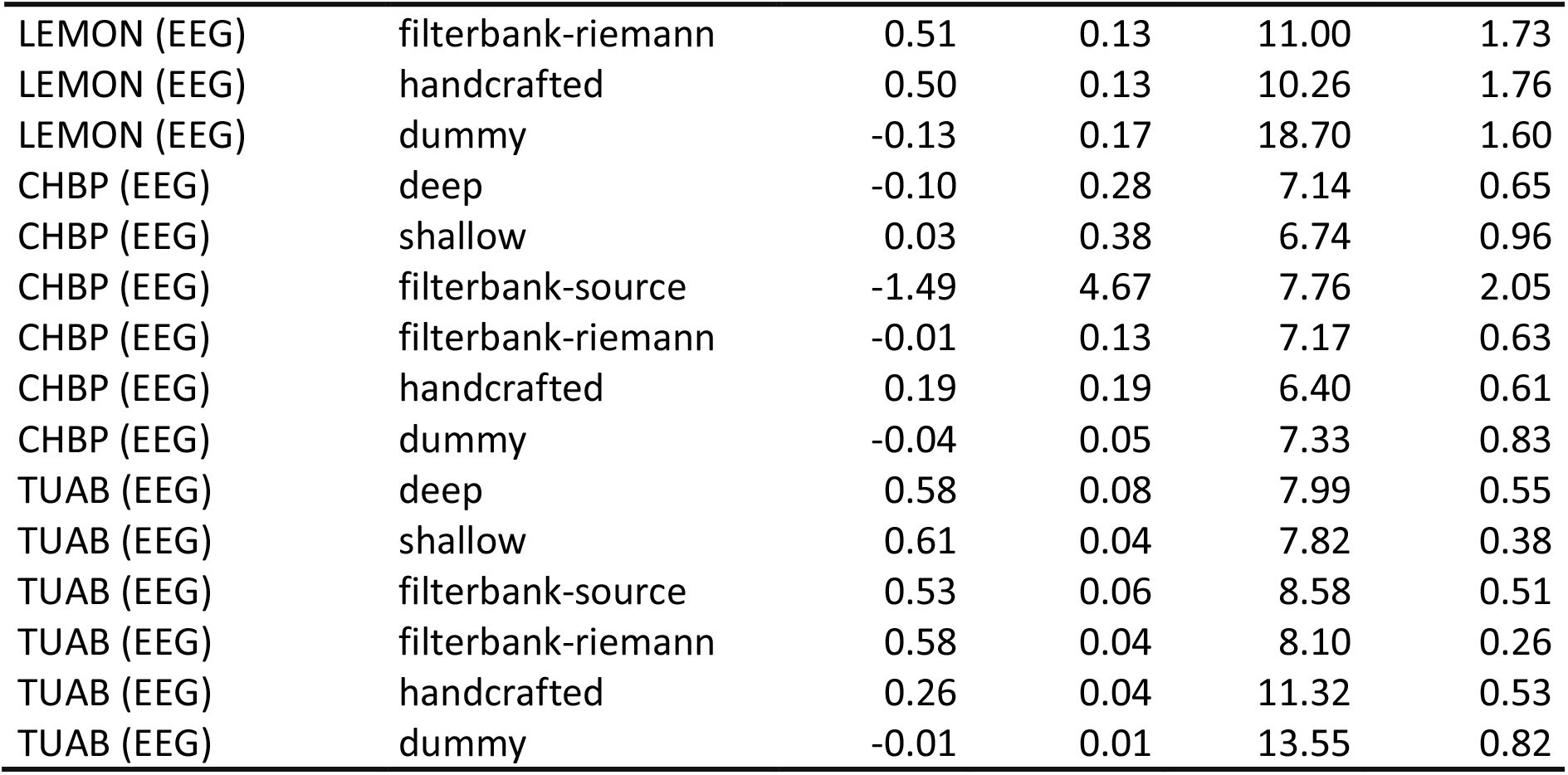
Aggregate cross-validation results across benchmarks and datasets.

| dataset | benchmark | $R^2_{(M)}$ | $R^2_{(SD)}$ | $MAE_{(M)}$ | $MAE_{(SD)}$ |
| --- | --- | --- | --- | --- | --- |
| Cam-CAN (MEG) | deep | 0.66 | 0.05 | 8.29 | 0.74 |
| Cam-CAN (MEG) | shallow | 0.69 | 0.03 | 8.14 | 0.90 |
| Cam-CAN (MEG) | filterbank-source | 0.69 | 0.06 | 8.10 | 1.11 |
| Cam-CAN (MEG) | filterbank-riemann | 0.72 | 0.05 | 7.65 | 0.81 |
| Cam-CAN (MEG) | handcrafted | 0.49 | 0.07 | 10.65 | 0.98 |
| Cam-CAN (MEG) | dummy | -0.02 | 0.03 | 15.90 | 1.22 |
| LEMON (EEG) | deep | 0.65 | 0.20 | 7.78 | 2.25 |
| LEMON (EEG) | shallow | 0.69 | 0.08 | 8.80 | 1.58 |
| LEMON (EEG) | filterbank-source | 0.65 | 0.12 | 8.93 | 1.56 |
| LEMON (EEG) | filterbank-riemann | 0.51 | 0.13 | 11.00 | 1.73 |
| LEMON (EEG) | handcrafted | 0.50 | 0.13 | 10.26 | 1.76 |
| LEMON (EEG) | dummy | -0.13 | 0.17 | 18.70 | 1.60 |
| CHBP (EEG) | deep | -0.10 | 0.28 | 7.14 | 0.65 |
| CHBP (EEG) | shallow | 0.03 | 0.38 | 6.74 | 0.96 |
| CHBP (EEG) | filterbank-source | -1.49 | 4.67 | 7.76 | 2.05 |
| CHBP (EEG) | filterbank-riemann | -0.01 | 0.13 | 7.17 | 0.63 |
| CHBP (EEG) | handcrafted | 0.19 | 0.19 | 6.40 | 0.61 |
| CHBP (EEG) | dummy | -0.04 | 0.05 | 7.33 | 0.83 |
| TUAB (EEG) | deep | 0.58 | 0.08 | 7.99 | 0.55 |
| TUAB (EEG) | shallow | 0.61 | 0.04 | 7.82 | 0.38 |
| TUAB (EEG) | filterbank-source | 0.53 | 0.06 | 8.58 | 0.51 |
| TUAB (EEG) | filterbank-riemann | 0.58 | 0.04 | 8.10 | 0.26 |
| TUAB (EEG) | handcrafted | 0.26 | 0.04 | 11.32 | 0.53 |
| TUAB (EEG) | dummy | -0.01 | 0.01 | 13.55 | 0.82 |

Interestingly, none of these approaches involving spatially fine-grained representations of the M/EEG signal worked well on the CHBP data, whereas the random forest on top of hand-crafted features scored systematically better than the dummy baseline. This may be related to three factors that come together in the CHBP benchmark dataset: Like the LEMON dataset, the sample size is relatively small. Second, the age distribution is far less uniform, leading to underrepresentation of elderly participants. This makes the learning task at hand harder as models have fewer training examples from elderly populations. These challenges apply equally to all machine learning benchmarks, hence, do not explain why the random forest model on hand-crafted features is working to some extent. In this context, it may be worthwhile to consider that the CHBP uses two different EEG montages, one with 60, one with 120 electrodes, which may induce strong difference in the covariance structure of the signals due to montage-specific noise structure related to the number of electrodes. This may have affected the random-forest pipeline less strongly as the hand-crafted features extracted marginal channel-wise summary statistics of the time-series or the power spectrum rather than pairwise interactions. Progress on this specific benchmark may therefore involve explicit consideration of the montage when selecting samples for cross-validation or even at the level of the machine learning model (e.g., by including the number of electrodes or montage type as covariate). Moreover, future availability of samples from older populations in the CHBP dataset will help disambiguate this point. Finally, once the expert-based quality-control annotations are considered for epochs-selection, the results obtained in this benchmark may change (see section Datasets/CHBP EEG data for details).

A different type of challenge is illustrated by the benchmarks on the LEMON dataset. As the age distribution is bimodal here (Fig. 2), the R^2^ score is not well calibrated as the average predictor will not provide a reasonable summary of the distribution. This is not automatically mitigated by considering the MAE as a metric. On the other hand, it will not affect the ranking of the machine learning models, which compare overall well to results obtained on the Cam-CAN and the TUAB datasets. To obtain a more rigorous baseline, one could envision a group-wise average predictor that, depending on the age group, would return the groups’ respective average age from the training data. We did not implement such a custom baseline here as it was our goal to stick to standard routines provided by the software libraries our benchmarks were based on. Second, it was our intention to expose such issues as this may stimulate future research and development.

## Discussion

In this study, we proposed empirical benchmarks for age prediction comparing distinct machine learning approaches across diverse M/EEG datasets comprising, in total, more than 2500 recordings. The benchmarks were implemented in Python based on the MNE-software ecosystem, the Braindecode package and the BIDS data standard. The explicit reliance on the BIDS standard renders these pipelines applicable to any M/EEG data presented in the BIDS format. This enabled coherent side-by-side comparisons of classical machine learning models and deep learning methods across M/EEG datasets recorded in different research or medical contexts.

Our cross-dataset and cross-model benchmarks pointed out stable ranking of model performance across two metrics, the R^2^ score and mean absolute error (MAE). R^2^ scores have been less consistently reported in the literature, however, the top MAE scores observed across datasets in this benchmark of 7 to 8 years are well in line with reports from previous publications (Sun et al. 2019; Sabbagh et al. 2020; Xifra-Porxas et al. 2021). While direct comparisons against MRI were not performed in this study, the present benchmarks would be compatible with the impression that for what concerns the overall performance of age prediction, M/EEG features are slightly weaker than MRI features (Engemann et al. 2020; Xifra-Porxas et al. 2021). We found that, overall, Riemannian filterbank models and deep learning models achieved the highest scores, whereas random forests based on hand-crafted features delivered robust performance. In line with previous work (Gemein et al. 2020), these results suggest that deep learning methods do not necessarily show a consistent advantage over classical pipeline models: Similar performance may be explained by the fact that our filterbank models and the deep models imply similar spatially aware representations of the M/EEG data (see results section for detailed discussion in context). Moreover, given the relatively small training datasets, it can be considered good news that these parameter-rich models did not seem to overfit as was evidenced by comparisons against simpler classical models. Yet, it may be simply a matter of collecting larger samples until deep learning approaches may reveal their advantage at extracting more elaborate representations of M/EEG signals. This may lead to positioning M/EEG-based brain age prediction on par with MRI-based brain age prediction just as MRI-based deep learning models of brain age have defined state-of-the-art performance on large datasets (Cole et al. 2017; Bashyam et al. 2020; Jonsson et al. 2019). However, more importantly, the value of M/EEG-derived brain age models should not be defined in terms of incremental improvement over MRI-based models as M/EEG-based models may enhance MRI-derived information (Engemann et al. 2020) or may be the only option available (Sun et al. 2019).

Our results nicely demonstrate a second critical merit of cross-model and cross-dataset benchmarking. It was sufficient to analyze four different sources of data until we found a perfectly legitimate EEG dataset from an academic research context (CHBP) in which our previously favored modeling techniques developed on the Cam-CAN and the TUAB data did not perform well by default. There may be good reasons for these discrepancies related to the age distribution found in the CHBP data and the fact that multiple different EEG montages were used in that dataset (see results section for detailed discussion in context). But more importantly, we did not anticipate this to happen and would have never learned about it had we confined the scope to previously analyzed datasets. Such discoveries are favored by systematic benchmarks with dataset-independent code implementation, which has the potential to lower the burden threshold for including always more datasets into model development. In the long run, we hope that this effort will stimulate new research leading to more generalizable models.

This brings us to some limitations of this work. Our work has been motivated by the absolute necessity to diversify datasets for development of M/EEG-based measures of brain health. This has led us to analyzing more than 2500 M/EEG recordings and, yet we only included four datasets. Other M/EEG datasets come to mind that would have been potentially relevant. The Human Connectome Project MEG data (Larson-Prior et al. 2013) includes MEG recordings from less than 100 participants, which we deemed insufficient for predictive regression modeling. The OMEGA data resource (Niso et al. 2016) was not accessible at the time of this investigation but would have been a good match for this study. Finally, the LIFE cohort (Loeffler et al. 2015) includes a large number of EEG recordings of participants sampled from the general population yet follows a closed / controlled access scheme. The Healthy-Brain-Network EEG data (Alexander et al. 2017) concerns a developmental cohort. Despite potentially relevant similarities between brain development and aging, age prediction in developmental cohorts would have exceeded the scope of the present study. Even if we had integrated these resources in the present benchmark, this may have only marginally enhanced the diversity covered by the current selection datasets as most public neuroscience datasets come from the wealthiest nations. We hope that this situation will improve as new promising international consortia and efforts emerge that focus on curating large EEG datasets from diverse national and cultural contexts (Ibanez et al. 2021; Shekh Ibrahim et al. 2020; “Global Brain Consortium Homepage”). A second limitation of the present study concerns the depth of validation. To advance our understanding of M/EEG-derived brain age, more systematic comparisons against MRI-derived brain age (Xifra-Porxas et al. 2021) and other measures of mental health and cognitive function are important objectives (Anatürk et al. 2021; Dadi et al. 2021).

In the following we wish to point out a few imminent opportunities for turning the limitations of the present work into future research projects, potentially, enabled by the results and tools brought by the current benchmarks.

### Opportunities and suggestions for follow-up research using the benchmark tools

#### Model averaging

In many instances, combining prediction models using model averaging approaches can improve the prediction performance (O’Connor et al. 2021; Dadi et al. 2021; Varoquaux et al. 2017). This could also be a practical way of combining the benchmarks into a single model for subsequent generalization testing. Future studies could use this benchmark to investigate model averaging approaches.

#### The impact of deeper architectures

An important design decision in deep neural networks is the total depth of the neural network. Here we used previously published architectures designed for EEG-based pathology decoding (Schirrmeister et al. 2017). Future studies could build on top of this benchmark to explore the importance of deep architectures for brain age modeling. Specifically, it would be possible to use methods from neural architecture search, e.g., AutoPyTorch (Zimmer, Lindauer, and Hutter 2021), to design better-performing architectures. Since this benchmark does not only provide access to diverse datasets in an identical file format, but also enables direct comparison to others, it is the optimal starting point for such an optimization while at the same time avoiding overfitting the architecture to a single dataset.

#### The role of preprocessing

While data cleaning is of major importance for extracting physiologically interpretable biomarkers, predictions from machine learning models tend to be far less affected by noise (Sabbagh et al. 2020). On the other hand, artifacts and noise may inform predictions, potentially reducing biological specificity. Future studies could benefit from this benchmark to quantify the role of artifact signals for brain age predictions and develop de-confounding strategies (Du et al. 2021; Mehrabi et al. 2021; Lu, Schölkopf, and Hernández-Lobato 2018; Bica, Alaa, and Van Der Schaar 2020).

#### Eyes-open versus eyes-closed

Some of the datasets analyzed in this benchmark contain resting-state signals under different conditions. In the lack of strong a-priori hypotheses, here we simply pooled both conditions. It is currently unclear whether the relationship between eyes-closed versus eyes-open resting-state may contain valuable information about brain aging. It is imaginable, however, that signals induced by transient visual deprivation may reveal levels of vigilance (Wong, DeYoung, and Liu 2016), which in turn may be altered by neuropsychiatric conditions (Hegerl et al. 2012). Future work could benefit from the benchmark to investigate the importance of eyes-closed versus eyes-open resting-state for brain age modeling.

#### Model inspection

The interpretability of machine learning models is essential for clinical impact (Rudin 2019; Ghassemi, Oakden-Rayner, and Beam 2021). This benchmark did not cover methods for explaining the role of variable importance for model predictions. Future work could validate the relative importance of M/EEG signals or features for brain age modeling.

#### Exploring the link with MRI and cognitive scores

This study established the tools and methods for basic benchmarks on prediction performance. However, to build useful brain age models, it is essential to validate brain-age predictions to cognitive function, measures of health or clinical endpoints (Dadi et al. 2021; Cole et al. 2018; Liem et al. 2017). To further establish the relative merit of M/EEG over MRI, comparisons between the modalities are essential (Engemann et al. 2020). Most of the datasets covered in this benchmark include MRI data, social details and psychometric scores next to the M/EEG data, providing a wealth of opportunities for deep cross-dataset validation of brain age measures.

## Conclusion

Computational benchmarks across M/EEG datasets and machine learning methods bear the potential to enhance applications of machine learning in clinical neuroscience in several ways. Standardization of data formats, software and analysis pipelines are important factors for the scalability of predictive modeling of M/EEG. For stimulating the development of more generalizable machine learning models it is crucial that a critical mass of M/EEG datasets be analyzed by the international community. As the diversity of the datasets increases, generalization gaps will manifest themselves, calling for computation methods for closing these gaps. The implied learning process may eventually lead to developing more widely applicable M/EEG-based biomarkers that are clinically robust across a wide range of sociocultural contexts, clinical populations, recording sites and measurement techniques. We hope that benchmarks, tools and resources resulting from this study will facilitate investigating open scientific questions related to learning biomarkers of brain health on an ever-growing number of M/EEG datasets from increasingly diverse real-world contexts.

## Acknowledgements

We would like to thank Pedro Valdés-Sosa and Jorge Bosch-Bayard and their team for the support with the Cuban Human Brain Project data and for the stimulating discussion. We would also like to thank Joseph Picone for the support with Temple University Hospital data. Finally, we would like to thank Anahit Babyan, Vadim Nikulin and Arno Villringer for the support with the LEMON dataset and the stimulating scientific discussions. We would like to thank Rik Henson and Ethan Knights for providing the Can-CAN datasets in BIDS format. We also thank Ana Radanovic and the other participants of the IDESSAI 2021 Inria-DFKI summer school for feedback and testing on earlier versions of the code and analysis developed in this project. This work was supported by the ANR BrAIN AI chair (ANR-20-CHIA-0016) and ANR AI-Cog (ANR-20-IADJ-0002) as well as by the Freiburg Graduate School of Robotics.

## Author contributions

authors in alphabetical order

**Conceptualization**: A.G., D.E.

**Data curation**: A.G., A.M., D.E., H.B., L.G.

**Software**: A.G., A.M., D.E., D.S., H.B., L.G., R.H.

**Formal analysis**: A.G., D.E.

**Supervision**: A.G., D.E.

**Funding acquisition**: A.G., D.E.

**Validation**: A.G., D.E.

**Investigation**: A.G., A.M., D.E., H.B., L.G.

**Visualization**: A.P., D.E.

**Methodology**: A.G., D.E.

**Project administration**: D.E.

**Writing—original draft**: D.E.

**Writing—review and editing**: A.G., A.M., D.E., D.S., H.B., L.G., T.B.

## Declaration of conflicts of interest

D.E. is a full-time employee of F. Hoffmann-La Roche Ltd.

H.B. receives graduate funding support from InteraXon Inc.

## Footnotes

1 https://github.com/meeg-ml-benchmarks/meeg-brain-age-benchmark-paper

2 https://github.com/coffeine-labs/coffeine

3 https://braindecode.org

4 https://github.com/mne-tools/mne-bids-pipeline

5 https://github.com/meeg-ml-benchmarks/meeg-brain-age-benchmark-paper

